# Physiological Responses of The Leaves of a High-Altitude Plant *Picrorhiza kurroa* to Cold Stress

**DOI:** 10.1101/2023.05.21.541615

**Authors:** Shreya Agrawal, Pooja Saklani

## Abstract

There are various plants that grow in high elevations and experience various environmental stresses, cold stress being the most prevalent one. Plants undergo biochemical, metabolic, molecular, and physiological changes under cold stress hence, they adopt various mechanisms to tolerate it. The antioxidant defense system, osmotic regulators and photosynthetic pigments in the plant provide them stress tolerance. The present study is conducted on a high-altitude plant, *Picrorhiza kurroa* which grows in such environmental conditions, to study the physiological parameters that provides it the coping mechanism against the cold stress. For this study, the leaves were collected from Pothivasa (2200 m asl) and Tungnath (3600 m asl) in Rudraprayag, Uttarakhand, India. The photosynthetic pigments, lipid peroxidation, antioxidant enzymes, and osmoprotectants present in the leaves were determined to visualize the impact of cold stress. It was revealed that the concentration of chlorophyll a, chlorophyll b, and carotenoids increased with increase in elevation. The activity of SOD, CAT, POD, APOX and GR were analyzed and they were observed to decrease with altitude. The MDA concentration which is an indicator of lipid peroxidation is higher in Pothivasa (2.214 μmol L^-1^) and lower in Tungnath (1.697 μmol L^-1^). There is a significant increase in the total protein content, total soluble sugar content and total proline content along the altitudinal gradient. Therefore, the leaves from both the sampling locations revealed the physiological changes occurred in them to adapt to the cold stress conditions.

## 1. INTRODUCTION

Stress is defined as “Any substance or condition which stops a plant’s metabolism, development or growth” **(Lichtenthaler, 2006)**. Both biotic and abiotic factors have an impact on plant productivity and growth **(Seki et al., 2003)**. Abiotic stress factors include water deficit, salinity, light stress, chilling, heat, flooding and soil compaction, freezing, trace elements toxicity, and mineral nutrient deficiencies. In order to maintain growth and developmental processes, as well as cellular homeostasis under such challenging circumstances, plants undergo biochemical, metabolic, molecular, and physiological changes when the ideal ambient temperature varies. When the temperature is below the optimum level, then the plants are said to be under cold stress. If the temperature is between 0-15°C, then it is termed as chilling stress **(Thomashow, 1999; Cook et al.,2004)**. In most hilly areas, cold stress is a prominent abiotic stress which hampers the normal growth of the plant. The cold stress causes dehydration, growth inhibition, metabolite imbalance, metabolic malfunction, plasma membrane rupture, and solute leakage in plants, all of which can result in plant mortality (**Mittler, 2002)**. On the other hand, exposure to a lower temperature can induce cold stress tolerance in plants. This phenomenon expresses various biochemical, metabolic, and molecular processes (**Zhu et al., 2007**). On exposure of plants to stress related to temperature, there is a modification of the metabolism of plants in two steps. Initially, by trying to alter their cellular metabolism, which is influenced by temperature fluctuations. Then, by modification of metabolism by enhanced tolerance mechanisms **(Schwender et al., 2004; Fernie et al., 2005)**.

Cold stress induces oxidative damage in the plant, causing the overaccumulation of reactive oxygen species (ROS), viz., hydrogen peroxide, hydroxyl ions, singlet oxygen, and Superoxide anion. Antioxidant enzymes play a vital role in inhibiting lipid peroxidation caused by ROS. Antioxidant enzymes include ascorbate oxidase (EC 1.10. 3.3), ascorbate peroxidase (APX; EC 1.11.1.1), catalase (CAT; EC 1.11.1.6), glutathione peroxidase (GPX; EC 1.11.1.9), glutathione reductase (EC 1.6.4.2), guaiacol peroxide (EC 1.11. 1.7), polyphenol oxidase (EC1.10.3.1), superoxide dismutase (SOD; EC 1.15.1.1), and so on.

Lipid peroxidation causes membrane damage in plants which indicates the cold stress and the malondialdehyde (MDA) is the indicator of lipid peroxidation in plants (**Lyons, 1973**). Many metabolites can function as osmoprotectants that induce tolerance during stress conditions (**Levitt, 1972; Guy, 1990; Nayyar et al., 2005; Farooq et al., 2009)**. They regulate the water balance within cells and lessen cellular dehydration. In addition, they can stabilize enzymes, membranes, and other biological components because of the behavior of their compatible solutes. These osmoprotectants include amino acids, lipids, organic acids, polyamines, and soluble carbohydrates (**Yadav, 2010**).

There is also a visible change in the morphology of the leaves like reduction in the length and width, and increment in its thickness in response to high elevation (**Roderick et al., 2000; Kok and Bahar, 2015**). The study of the leaves of a plant grown in the Northwestern Himalayas makes it ideal to study the cold stress tolerance and adaptations. Therefore, in order to investigate the physiological factors influencing this plant’s ability to withstand cold stress, the current study is being conducted on *Picrorhiza kurroa* Royle ex Benth. (Kutki), which belongs to the family Scrophulariaceae and is distributed in the sub-alpine and alpine regions of the North Western Himalayas between 3000-5000 m asl (**Chettri et al., 2005**). The Western Himalayan states of Jammu and Kashmir, Himachal Pradesh, and Uttarakhand are home to *Picrorhiza kurroa* (**Kaul et al., 1996**). It has a mild, unpleasant aroma and a bitter flavor. The iridoid glycosides secreted by this plant infer implicit medicinal values to it. These glycosides include active substances such as apocynin, drosin, and cucurbitacins, as well as the picrosides I, II, III, and V and kutkosides (**Jia et al., 1999; Stuppner et al., 1989**). The plant has excellent economic worth due to its ability to treat various illnesses, including leishmaniasis, anemia, jaundice, inflammation, and gastrointestinal problems (**Ghisalberti, 1998; Sharma et al., 2011**).

## 2. MATERIALS AND METHODS

The biological replicates of the leaves of *Picrorhiza kurroa* were collected in the month of July from the nurseries of HAPPRC, HNBGU, located in Pothivasa (30°28′N lat; 79°16′E long at an altitude of 2200 m asl) and Tungnath (30°14′N Lat; 79°13′E Long at an altitude of 3600 m asl) in Rudraprayag district, Uttarakhand, India (Fig. 1 & 2). There is an altitudinal variation of about 1400 m in both the sampling locations. The temperatures in Pothivasa and Tungnath sampling sites observed during sampling were 13°C and 1°C, respectively.

**Fig. 1:**
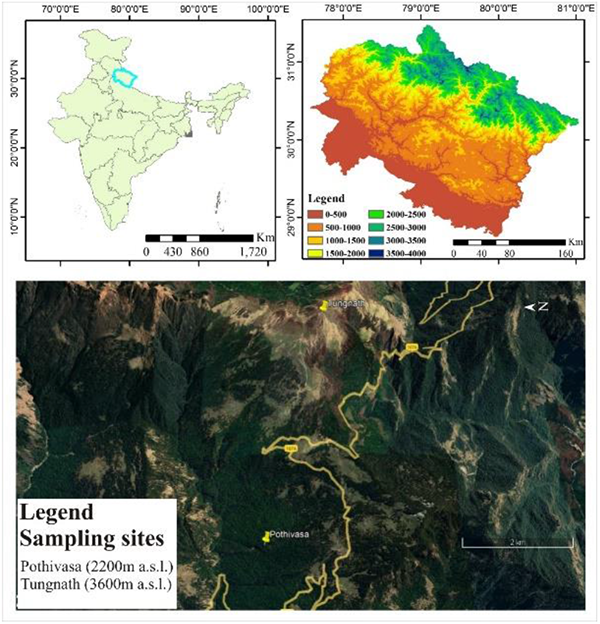
Location map of the study area Pothivasa (2200m asl) and Tungnath (3600m asl)

**Fig. 2:**
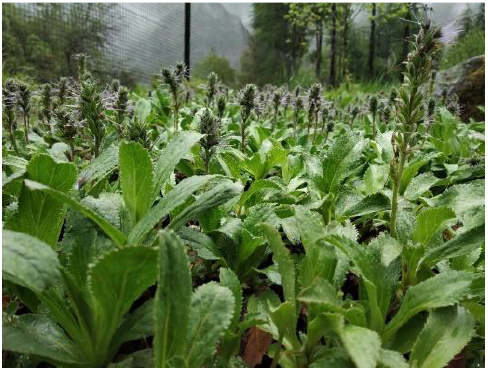
*Picrorhiza kurroa* cultivated in Pothivasa

### 2.1. PHOTOSYNTHETIC PIGMENTS

The fresh leaves of *Picrorhiza kurroa* from Pothivasa (PVLE) and Tungnath (TNLE) were grounded with ethanol, extracted completely and then filtered to determine the photosynthetic pigments in the leaves. The absorbance of chlorophyll a (Chl a), chlorophyll b (Chl b), and total carotenoids (Car) were all measured at wavelengths of 665 nm, 649 nm, and 470 nm, respectively. Their concentrations (mg g^-1^ DW) were then determined as per Wellburn and Liehtenthaler (1983) using following equations:

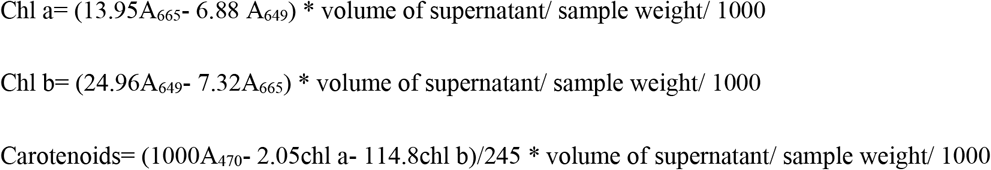

### 2.2. ANTIOXIDANT ENZYMES ACTIVITY

In ice-cold conditions, 5 gm of leaves were homogenised in 0.15 M potassium phosphate buffer at pH 7.0 before being centrifuged at 15000 rpm for 20 min. For the investigation of enzymatic antioxidant activity and protein estimation, the supernatant, or enzyme extract, was collected and stored at -20°C.

#### 2.2.1. Superoxide Dismutase (SOD) (EC. 1.15.1.1)

The activity of SOD was determined according to the method by Kono (1978). The reaction mixture was prepared using sodium carbonate buffer, NBT, Triton X-100, hydroxylamine hydrochloride and enzyme extract. The amount by which the rate of NBT degradation was inhibited was determined by a rise in absorbance at 540 nm. The percent inhibition of NBT reduction (y%) and sample producing 50% inhibition (z μl) were calculated.

#### 2.2.2. Catalase (Cat) (EC. 1.11.1.6)

According to Aebi (1983), phosphate buffer, hydrogen peroxide, and enzyme extract were added to formulate the reaction mixture. At 240 nm, the absorbance was measured. The rate of decomposition of H_2_O_2_ decreased the absorbance. Utilizing the molar Extinction coefficient of 6.93*10^−3^ mM^-1^ cm^-1^, the enzyme activity was computed.

#### 2.2.3. Guaiacol Peroxidase (POD) (EC. 1.11.1.7)

According to Putter (1974), to determine the activity of POD enzymes in leaves, a test cuvette was filled with phosphate buffer, guaiacol solution, enzyme sample, and H_2_O_2_ solution. At 436 nm, the rate of GDHP production was monitored spectrophotometrically. When calculating enzyme activity, the extinction coefficient was taken at 25 mM^-1^ cm^-1^.

#### 2.2.4. Ascorbate Peroxidase (APOX) (EC. 1.11.1.11)

Using 3 ml of the reaction mixture, which included phosphate buffer, ascorbate, H_2_O_2_, and enzyme extract, the decrease in absorbance was measured at 290 nm as per Nakano and Asada (1981). The enzyme activity was calculated using the extinction coefficient as 2.8 mM^-1^ cm^-1^.

#### 2.2.5. Glutathione Reductase (GR) (EC. 1.6.4.2)

GR activity was determined according to Carlberg and Mannervik (1975) by detecting the oxidation of NADPH at 340 nm in a reaction mixture containing phosphate buffer, EDTA, NADPH, and oxidised glutathione (GSSG). The minute decrease in absorbance was followed at 340 nm. The enzyme activity was calculated using the extinction coefficient (6.22 mM^-1^cm^-1^).

### 2.3. OSMOPROTECTANTS

#### 2.3.1. Total Soluble Sugar

The determination of Total Soluble Sugar content was done according to Hedge and Hofreiter (1962). The sample was hydrolyzed with 2.5 N HCl for three hours in water bath at 100°C, and then cooled to room temperature. Once the effervescence had stopped, it was neutralised using sodium carbonate. The volume was then increased to 100 ml, centrifuged, and the supernatant was collected and used for analysis. Working standards were also prepared and anthrone reagent was added in all the test tubes. The mixture was first heated in boiling water and then cooled quickly and the absorbance was taken at 630 nm.

#### 2.3.2. Total Protein

Lowry’s method was employed for the determination of protein content. The reagents and protein standard stock solution containing BSA were prepared and mixed. They underwent a 30-minute incubation period at ambient temperature and in complete darkness. Blue color was developed. At 550 nm, the readings were recorded. The amount of protein in the samples was determined by plotting an absorbance vs. concentration graph for standard solutions of proteins. Protein content was expressed as mg/g FW.

#### 2.3.3. Total Proline

Total Proline content was estimated according to the protocols by Bates *et al*. (1973). The frozen leaves were mixed with sulphosalicylic acid to homogenise it. The residue was centrifuged at 12000 rpm for 10 minutes to separate it. The process was stopped in an ice bath after the dissolving equal amounts of acid-ninhydrin, glacial acetic acid and homogenised tissue. 2 ml of toluene was used to extract the reaction mixture, which was then violently agitated and kept at room temperature to allow the two phases to separate. The absorbance of 1 ml of upper phase and toluene (blank) was taken at 520 nm. D-Proline was used to prepare a standard curve to calculate the proline concentration.

### 2.4. DETERMINATION OF LIPID PEROXIDATION

The leaves were ground in trichloroacetic acid (TCA), centrifuged and then supernatant was collected and mixed with Thiobarbituric Acid (TBA). It was heated and then cooled in an ice bath. It was centrifuged again and the absorbance of the supernatant was determined at wavelengths of 450 nm, 532 nm, and 600 nm. The malondialdehyde (MDA) content was computed using the following equation and represent it as μmolg^-1^ dry weight:

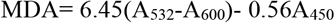

## 3. RESULTS

### 3.1. Photosynthetic pigments

Effect of altitudinal variation was studied on chlorophyll a, chlorophyll b and carotenoids present in the leaves of *Picrorhiza kurroa* which were collected from two locations i.e., Pothivasa and Tungnath, at different elevations. It was observed that the concentration of chlorophyll a, chlorophyll b and carotenoids in PVLE were 0.072 mg g^-1^ DW, 0.29 mg g^-1^ DW and 0.065 mg g^-1^ DW, respectively and that in TNLE were 0.075 mg g^-1^ DW, 0.304 mg g^-1^ DW, and 0.072 mg g^-1^ DW, respectively. Hence, these pigments showed positive correlation with the altitude i.e., TNLE contained higher levels of photosynthetic pigments than PVLE (Fig.3).

**Fig. 3:**
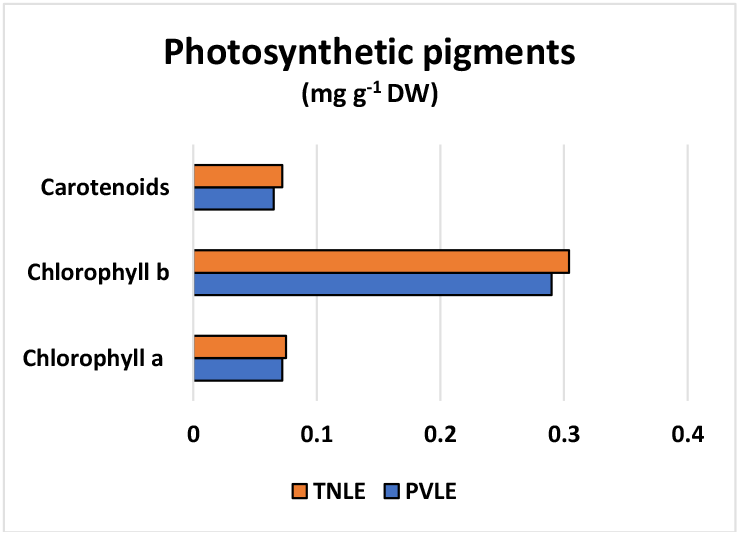
Photosynthetic pigments (Chl a, Chl b and carotenoids) present in the leaves extract

### 3.2. Enzymatic Antioxidants

The activity of antioxidant enzymes is affected in high altitudes. Superoxide dismutase, catalase, guaiacol peroxidase, ascorbate peroxidase and glutathione reductase were studied for their change in activity with respect to the altitude. SOD results revealed that the sample required for 50% inhibition of NBT reduction was very high in case of PVLE (286± 13.93 μl) in comparison to TNLE (4.82±0.61 μl) (Table 1). Other enzymes viz., Cat, POD, APOX and GR also showed the higher specific activity in PVLE than TNLE (Fig. 4). The specific activity of Cat, POD, APOX and GR in PVLE were calculated to be 20.41± 0.407 U/mg protein/min, 14.42± 0.67 U/mg protein/min. 3.34± 0.25 U/mg protein/min, and 0.17 ± 0.01 U/mg protein/min, respectively and that in TNLE were 7.01± 0.99 U/mg protein/min, 4.10 ± 0.13 U/mg protein/min, 1.63± 0.21 U/mg protein/min, and 0.77± 0.01 U/mg protein/min, respectively. Hence, all these enzymes showed negative correlation with respect to the altitude.

**Table 1:** Activity of SOD enzyme; y: % inhibition of NBT and z: μl of sample for 50% inhibition

| SOD | y (%) | z (μl) |
| --- | --- | --- |
| PVLE | $12.26 \pm 0.58$ | $286 \pm 13.93$ |
| TNLE | $733.61 \pm 86.08$ | $4.82 \pm 0.61$ |

**Fig. 4:**
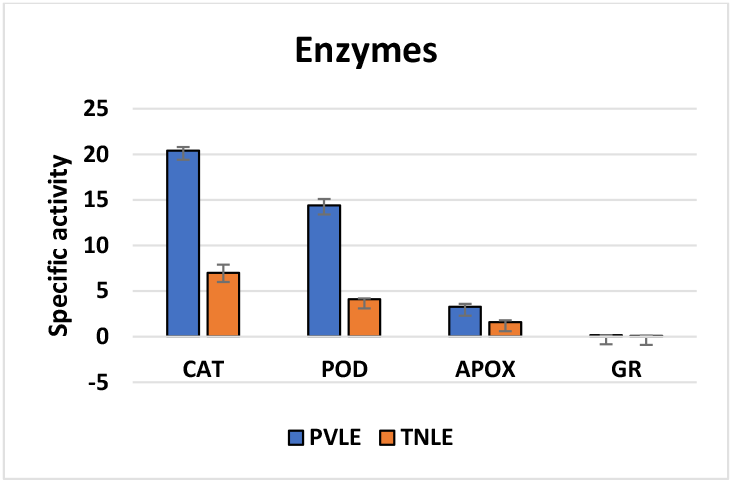
Specific activity of Cat, POD, APOX and GR enzymes

### 3.3. Osmoprotectants

Soluble protein, soluble carbohydrates and proline are the osmotic regulators which are accumulated or induced directly or indirectly by the abiotic stress in the environment **(Bartels et al., 2005)**. In the present study, it was observed that the concentration of these osmoprotectants increase with the increase in altitude as shown in Fig. 5. The total protein content was found to be 3.534 μg protein/100μl in PVLE and 4.806 μg protein/100μl in TNLE, Total soluble sugar was 0.35 mg sugar/ml in PVLE and 0.55 mg sugar/ml in TNLE, and Total proline content was 4.53 μmoles/gm and 6.73 μmoles/gm in PVLE and TNLE, respectively.

**Fig. 5:**
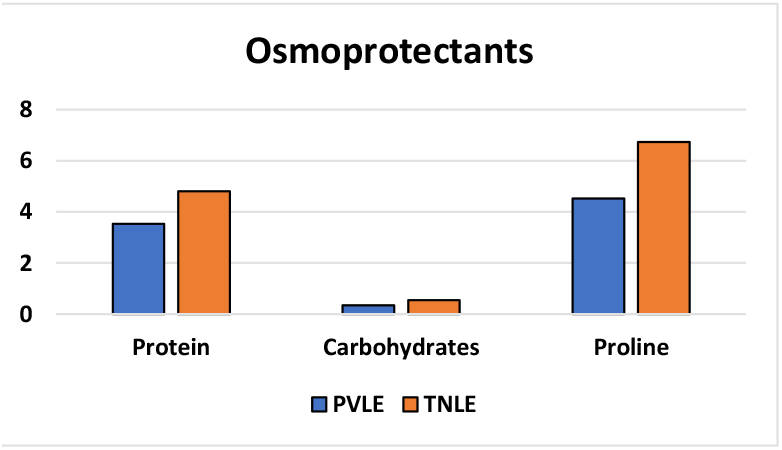
Concentration of soluble protein (μg protein/100μl), soluble carbohydrates (mg sugar/ml), and proline (μmoles/gm) in PVLE and TNLE

### 3.4. Lipid peroxidation

The oxidative damage was estimated by evaluating the Malondialdehyde (MDA) content, which acts as an indicator for lipid peroxidation. The result showed the increased levels of MDA in PVLE (2.214 μmol L^-1^) in comparison to TNLE (1.697 μmol L^-1^), representing negative correlation between the altitude and MDA concentration (Fig. 6).

**Fig. 6:**
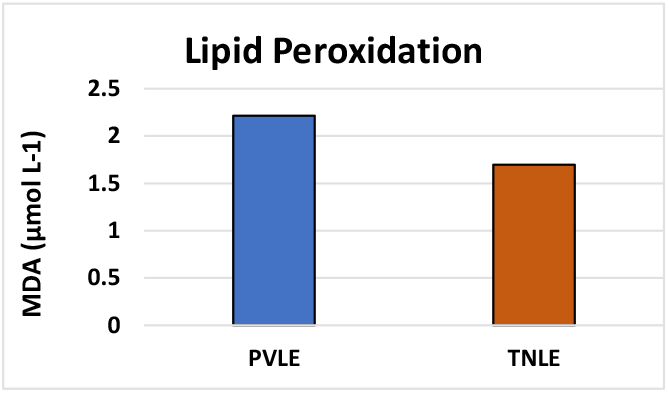
MDA concentration in leaves extract to analyze lipid peroxidation

## 4. DISCUSSION

The plant of the study, *Picrorhiza kurroa* is a high value medicinal herb that grows at high-altitude and faces the cold stress conditions. In high altitudes, there are various abiotic factors which includes light intensity, oxygen pressure, precipitation, photoperiod, temperature and UV intensity that act as stress and affects the plant’s growth (**Chen et al., 2014**). In general, different plant species copes up with the environmental stresses by employing different adaptation mechanisms such as altering their morphology, metabolism and physiology. Under stress, the life cycle of the plant is maintained by the antioxidant defense system, osmotic regulators and photosynthetic pigments (**Cui et al., 2016, Qin et al., 2005**). Plant may use all or some of these methods to combat with stress conditions.

In the current investigation, the concentration of chlorophyll present in the leaves of *P. kurroa* was seen to increase with altitude as chlorophyll a pigment increased from 0.072 mg g^-1^ DW to 0.075 mg g^-1^ DW, and chlorophyll b from 0.29 mg g^-1^ DW to 0.304 mg g^-1^ DW. Plant photosynthetic capacity is directly impacted by photosynthetic pigments, which are also vulnerable to environmental stresses and are responsible for assimilating and transforming light. Although, chlorophyll content can be decreased by water stress, strong light, and high temperature by slowing synthesis, increasing decomposition, or harming chloroplast structures (**Hazrati et al., 2016; Ashraf and Harris, 2013**). Other herbaceous species have also been shown to exhibit an upregulation of photosynthesis in response to cold stress, including *Malva neglecta* (**Verhoeven et al., 1999; Adams et al., 2001, 2002, 2004, and 2006**), winter cereals **(Hurry et al., 1995a, 1995b, 1995c**), and spinach (**Holaday et al., 1992; Adams et al., 1995; Martindale and Leegood, 1997**). This acclimatory elevation of photosynthetic capacity apparently implies an investment to increase the overall capacity of photosynthesis/size of the photosynthetic apparatus in order to compensate the decrease in enzyme activity at lower temperatures **(Adams et al., 2002)**. Carotenoids are the complexes which participate in photosynthesis and can remove the ROS which gets accumulated and causes photooxidation which damages the chloroplast of the leaves (**Armstrong and Hearst, 1996; Demmig-Adams, 1992**). In this study, carotenoids were observed to increase with the altitude, i.e., from 0.065 mg g^-1^ DW to 0.072 mg g^-1^ DW, which may be due to the pigments’ protective role in the dissipating the additional energy and ROS scavenging (**HM et al., 1999**). The similar pattern was observed in the skin tissues of peach (**Karagiannis, 2016**), even though the concentration of carotenoids generally decreases at high altitude.

The effect of altitudinal variation on the antioxidant enzymes like superoxide dismutase (SOD), catalase (CAT), guaiacol peroxidase (POD), ascorbate peroxidase (APOX), and glutathione reductase (GR) present in *P*.*kurroa* resulted in the decrease in their activity with the increase in altitude (Table 1 and Fig. 4). In high altitude and under stress conditions, a significant level of ROS causes oxidative damage to plants (**Asada, 1999**). Plants have evolved an antioxidant defense system that includes antioxidant enzymes in order to prevent oxidative damage under an array of difficult environmental conditions. The study of these enzymes on *Coleus forkohlii* along the altitudinal gradient showed the best enzymatic activity at highest altitude to combat oxidative stress **(Rana and Saklani, 2019)**. But, the results of this investigation suggesting the decrease in enzymatic activity, reveals that they were adequate to change O_2_ and H_2_O_2_ levels (**Wang et al., 2009**), which may suggest that *P. kurroa* is better able to adjust to altitudinal gradients.

Plants regulate the osmotic stress developed due to low temperature with the help of osmoprotectants. The cold stress causes the change in intracellular and metabolic pathways by synthesizing proteins, carbohydrates and proline (**Guy et al., 1992**). In this study, *P*.*kurroa* leaves also showed changes in response to the cold stress by increasing the synthesis of these osmoprotectants as shown in Fig. 5. These regulatory proteins controls and/ or regulates the expression and activity of genes related to stress, hence contributing in cold stress tolerance (**Mahajan and Tuteja, 2005**). The increase in carbohydrates helps in protecting the structure of cell membrane which plays an important role in plant stress tolerance (**Korner, 1999; Bano and Fatima, 2009**). The overproduction of proline also acts as cryoprotectant. Research report by **Kandler and Hopf (1982)** stated that there is an accumulation of sucrose under cold stress. **Bano and Fatima (2009)** also observed the higher concentration of sugar in high elevations. Researchers observed that the proline concentration significantly increase during cold hardening. (**Galiba et al., 1994; Mahajan and Tuteja, 2005**).

Reactive oxygen species (ROS) may accumulate up as a result of stress-related cellular damage, which can then result in membrane lipid peroxidation (**Demiral & Turkan, 2005**). Malondialdehyde (MDA), a byproduct of lipid peroxidation, has been widely employed as a marker of membrane damage brought on by free radicals under different abiotic conditions (**Alexieva et al., 2001**). Our research revealed that the MDA levels in *P. kurroa* leaves reduced from 2.214 μmol L^-1^ in PVLE to 1.697 μmol L^-1^ in TNLE, along the gradient of altitude. In the high-altitude region, less damage in plasma membrane and better ROS scavenging capabilities are may be responsible for the decline in MDA levels. The leaves of *L. secalinus* yielded - similar outcomes where in Minhe County (1872 to 2185 m asl) MDA content in the leaves initially elevated and then declined, and that in Huangzhong County (2163 to 2935 m asl) linearly declined along the altitudinal gradient. (**Cui et al., 2018**).

Therefore, the current study on the leaves of *Picrorhiza kurroa* showed the adaptation mechanism of this plant against the cold stress along the altitude. The increase in the concentration of chlorophyll pigments and carotenoids in the leaves showed their vital role in increasing the photosynthesis and scavenging the free radicals, respectively. The decrease in enzymatic antioxidant activity and lipid peroxidation reveals the adjustability of the leaves in high altitude. Since, the lipid peroxidation and antioxidant activity in the leaves is observed to be decreasing in TNLE, it may be assumed that the plant is not using the antioxidant mechanism to combat the cold stress, instead, the soluble sugar, protein and proline may be essential for osmotic regulation in high altitude according to the increase in their levels with elevation that has been seen in these plants. Hence, indicating the physiological changes in these leaves to tolerate the stress. Further studies can be done to learn about the molecular changes occurring in these leaves to survive the stress conditions.

## ACKNOWLEDGEMENT

The authors are thankful to The University Grants Commission (UGC), New Delhi for granting fellowship to the first author and the Department of Biotechnology, HNBGU for providing the facilities.

## REFERENCES

1. Adams WW III, Demmig-Adams B, Rosenstiel TN and Ebbert V (2001). Dependence of photosynthesis and energy dissipation activity upon growth form and light environment during the winter. Photosynthesis Research. 67: 51–62.

2. Adams WW III, Demmig-Adams B, Rosenstiel TN, Brightwell AK and Ebbert V (2002). Photosynthesis and photoprotection in overwintering plants. Plant Biology. 4: 545–557.

3. Adams WW III, Hoehn A and Demmig-Adams B (1995). Chilling temperatures and the xanthophyll cycle. A comparison of warm-grown and overwintering spinach. Australian Journal of Plant Physiology. 22: 75–85.

4. Adams WW III, Zarter CR, Ebbert V and Demmig-Adams, B (2004). Photoprotective strategies of overwintering evergreens. BioScience. 54: 41–49.

5. Adams WW III, Zarter CR, Mueh KE, Amiard V and Demmig-Adams B (2006). Energy dissipation and photoinhibition: a continuum of photoprotection. Dordrecht: Springer. 21: 49–64.

6. Aebi H (1984). Catalase in vitro. Methods Enzymol. 105: 121–126.

7. Alexieva V, Sergiev I, Mapelli S and Karanov E (2001). The effect of drought and ultraviolet radiation on growth and stress markers in pea and wheat. Plant, Cell & Environment. 24(12):1337–44.

8. Armstrong GA and Hearst JE (1996). Carotenoids 2: Genetics and molecular biology of carotenoid pigment biosynthesis. FASEB J. 10(2):228–37.

9. Asada K (1999). The water-water cycle in chloroplasts: Scavenging of Active Oxygens and Dissipation of Excess Photons. Annu Rev Plant Physiol Plant Mol Biol. 50(1):601–39.

10. Ashraf M and Harris PJC (2013). Photosynthesis under stressful environments: An overview. Photosynthetica. 51(2):163–90.

11. Bano A and Fatima M (2009). Salt tolerance in Zea mays (L.) following inoculation with Rhizobium and Pseudomonas. Biol. Fertility Soils. 45: 405–413.

12. Bartels D and Sunkar R (2005). Drought and salt tolerance in plants. Crit Rev Plant Sci. 24(1):23–58

13. Bates LS, Waldren RP and Teare ID (1973). Rapid determination of free proline for water-stress studies. Plant Soil. 39: 205–207. Berlin, Heidelberg, New York.

14. Carlberg INCER and Mannervik BENGT (1975). Purification and characterization of the flavoenzyme glutathione reductase from rat liver. Journal of biological chemistry, 250(14): 5475–5480.

15. Chen BX, Zhang XZ, Tao J, Wu JS, Wang JS and Shi PL (2014). The impact of climate change and anthropogenic activities on alpine grassland over the Qinghai-Tibet Plateau. Agr Forest Meteorol. 189(189):11–8.

16. Chettri N, Sharma E and Lama SD (2005). Non-timber forest produces utilization, distribution and status in a trekking corridor of Sikkim, India, Lyonia. 8:89–101.

17. Cook D, Fowler S, Fiehn O and Thomashow MF (2004). A prominent role for the CBF cold response pathway in configuring the low temperature metabolome of Arabidopsis, Proc. Natl Acad. Sci. (USA) 101, 15243–15248.

18. Cui G, Li B, He W, Yin X, Liu S, Lian L, Zhang Y, Liang W and Zhang P (2018). Physiological analysis of the effect of altitudinal gradients on Leymus secalinus on the Qinghai-Tibetan Plateau. PloS one. 13(9): e0202881.

19. Cui GX, Wei XH, Degen AA, Wei XX, Zhou JW and Ding LM (2016). Trolox-equivalent antioxidant capacity and composition of five alpine plant species growing at different elevations on the Qinghai-Tibetan Plateau. Plant Ecology & Diversity. 9(4):387–96.

20. Demiral T and Turkan I (2005). Comparative lipid peroxidation, antioxidant defense systems and proline content in roots of two rice cultivars differing in salt tolerance. Environ Exp Bot. 53(3):247–57.

21. Demmig-Adams B, Iii WWA (1992). Carotenoid composition in sun and shade leaves of plants with different life forms. Plant Cell & Environment. 15(4):411–9.

22. Farooq M, Wahid A, Kobayashi N, Fujita D and Basra SMA (2009). Plant drought stress: effects, mechanisms and management, Agron. Sustain. Dev. 29: 185–212.

23. Fernie AR, Geigenberger P and Stitt M (2005) Flux an important, but neglected, component of functional genomics, Curr. Opin. Plant Biol. 8: 174–182.

24. Galiba G, Sutka J, Snape JW, Tuberosa R, Quarrie SA, SarkadI L and Veisz O (1994). The association of forest resistance gene Frl with stress induced osmolyte and ABA accumulation in wheat. In: Proceedings of the workshop on crop adaptation to cool climates. Oct 12–14, Hamburg Germany, 389-401.

25. Ghisalberti EL (1998). Biological and pharmacological activity of naturally occurring iridoids and secoiridoids. Phytomed. 5: 147–163.

26. Guy CL (1990). Cold acclimation and freezing stress tolerance: role of protein metabolism, Annu. Rev. Plant Phys. 41: 187–223.

27. Guy CL, Huber JL and Huber SC (1992). Sucrose phosphate synthase and sucrose accumulation at low temperature. Plant Physiol. 100: 502–508.

28. H M and N KK (1999). The violaxanthin cycle protects plants from photooxidative damage by more than one mechanism. Proc Natl Acad Sci U S A. 96(15):8762–7.

29. Hazrati S, Tahmasebi-Sarvestani Z, Modarres-Sanavy SAM, Mokhtassi-Bidgoli A and Nicola S (2016). Effects of water stress and light intensity on chlorophyll fluorescence parameters and pigments of Aloe vera L. Plant Physiology and Biochemistry. 1ver06:141–8.

30. Hedge JE and Hofreiter BT (1962). Estimation of carbohydrate. Methods in carbohydrate chemistry. Academic Press, New York.17–22.

31. Holaday AS, Martindale W, Alred R, Brooks A and Leegood R (1992). Changes in activities of enzymes of carbon metabolism in leaves during exposure of plants to low temperature. Plant Physiology. 98: 1105–1114.

32. Hurry VM, Keerberg O, Pärnik T, Gardeström P and Öquist G (1995b). Cold-hardening results in increased activity of enzymes involved in carbon metabolism in leaves of winter rye (Secale cereale L.). Planta. 195: 554–562.

33. Hurry VM, Strand Å, Tobiæson M, Gardeström P and Öquist G (1995a). Cold hardening of spring and winter wheat and rape results in differential effects on growth, carbon metabolism and carbohydrate content. Plant Physiology. 109: 697–706.

34. Hurry VM, Tobiæson M, Krömer S, Gardeström P and Öquist, G (1995c). Mitochondria contribute to increased photosynthetic capacity of leaves of winter rye (Secale cereale L.) following cold-hardening. Plant, Cell and Environment. 18: 69–76.

35. Jia Q, Hong MF and Minter D (1999). Pikuroside: a novel iridoid from Picrorhiza kurrooa. J Nat Prod. 62:901–903.

36. Kandler O and Hopf H (1982). Oligo saccharides based on sucrose. In: Encyclopedia of plant Physiology. Berlin: Springer, 13A: 348–382.

37. Karagiannis E, Tanou G, Samiotaki M, Michailidis M, Diamantidis G and Minas IS (2016). Comparative Physiological and Proteomic Analysis Reveal Distinct Regulation of Peach Skin Quality Traits by Altitude. Frontiers in plant science. 7:1689.

38. Kaul MK and Kaul K (1996). Studies on medico-ethnobotany, diversity, domestication and utilization of Picrorhiza kurrooa, In: Supplement to cultivation and utilization of medicinal plants. Jammu, CSIR. 333–348.

39. Kok D and Bahar E (2015). Effects of different vineyard altitudes and grapevine directions on some leaf characteristics of cv. Gamay Vitis vinifera L. Bulg J Agric Sci. 21:320–324.

40. Kono Y (1978). Generation of superoxide radical during autoxidation of hydroxylamine and an assay for superoxide dismutase. Archives of biochemistry and biophysics. 186(1): 189–195.

41. Korner C (1999). Alpine plant life: functional plant ecology of high mountain ecosystems. Springer,

42. Levitt J (1972). Responses of Plants to Environmental Stresses, Academic Press, New York.

43. Lichtenthaler HK (2006). The stress concept in plants: an introduction. Ann NY Acad Sci. 851:187–198.

44. Lichtenthaler HK and Wellburn A (1983). Determination of total carotenoids and chlorophylls a and b of leaf in extracts in different solvents. Biochem. Soc. Trans. 603: 591–592.

45. Lowry OH, Rosebrough NJ, Farr AL and Randall RJ (1951). Protein measurement with the Folin phenol reagent. J. Biol. Chem. 193:265–275.

46. Lyons JM (1973). Chilling injury in plants. Annu. Rev. Plant Physiol. 24: 445–466.

47. Mahajan S and Tuteja N (2005). Cold, salinity and drought stresses: An overview. Arch. Biochem. Biophy. 444: 139–158.

48. Martindale W and Leegood RC (1997). Acclimation of photosynthesis to low temperature in Spinacia oleracea L. I. Effects of acclimation on CO_2_assimilation and carbon partitioning. Journal of Experimental Botany. 48: 1865–1872.

49. Mittler R (2002). Oxidative stress, antioxidants and stress tolerance. Trends in Plant Science. 7(9): 405–410.

50. Nakano Y and Asada K (1981). Hydrogen peroxide is scavenged by ascorbate-specific peroxidase in spinach chloroplasts. Plant and cell physiology. 22(5): 867–880.

51. Nayyar H, Chander K, Kumar S and Bains T (2005). Glycine betaine mitigates cold stress damage in Chickpea, Agron. Sustain. Dev. 25: 381–388.

52. Pütter J (1974). Peroxidases. In Methods of enzymatic analysis. Academic Press. 685–690.

53. Qin JH and Liu Q (2009). Impact of seasonally frozen soil on germinability and antioxidant enzyme activity of Picea asperata seeds. Can J For Res. 39(4):723–30.

54. Rana PS and Saklani P (2019). Analyzing effect of altitudinal variation in Enzymatic antioxidants of Coleus forskohlii from Uttarakhand, India. Plant Cell Biotechnology and Molecular Biology. 20(9&10):442–450.

55. Roderick ML, Berry SL, Noble IR (2000) A framework for understanding the relationship between environment and vegetation based on the surface area to volume ratio of leaves. Func Ecol 14:423–437.

56. Schwender J, Ohlrogge J and Shachar-Hill Y (2004) Understanding flux in plant metabolic networks. Curr. Opin. Plant Biol. 7: 309–317.

57. Seki M, Kamei A, Yamaguchi-Shinozaki K and Shinozaki K (2003). Molecular responses to drought, salinity and frost: common and different paths for plant protection. Curr Opin Biotechnol. 14:194–199.

58. Sharma PK, Thakur SK, Manuja S, Rana RK and Kumar P (2011). Observations on traditional phytotherapy among the inhabitants of Lahaul Valley through Amchi system of medicine—A cold desert area of Himachal Pradesh in North Western Himalayas, India. Chi Med. 2: 93–102.

59. Stuppner H and Wagner H (1989). New cucurbitacin glycosides from Picrorhiza kurrooa. Plant Med. 55:559–563.

60. Thomashow M.F. (1999) Plant cold acclimation: freezing tolerance genes and regulatory mechanisms, Annu. Rev. Plant Phys. 50, 571–599.

61. Verhoeven AS, Adams WW III and Demmig-Adams B (1999). The xanthophyll cycle and acclimation of Pinus ponderosa and Malva neglecta to winter stress. Oecologia. 118: 277–287.

62. Wang Y, He W, Huang H, An L, Wang D and Zhang F (2009). Antioxidative responses to different altitudes in leaves of alpine plant Polygonum viviparum in summer. Acta Physiologiae Plantarum. 31(4):839–48.

63. Yadav SK (2010). Cold stress tolerance mechanisms in plants. A review. Agronomy for sustainable development. 30(3): 515–527.

64. Zhu J, Dong CH and Zhu JK (2007). Interplay between cold-responsive gene regulation, metabolism and RNA processing during plant cold acclimation, Curr. Opin. Plant Biol. 10: 290–295.

